# Plastic loss of motile cilia in the gills of *Polypterus* in response to high CO_2_ or terrestrial environments

**DOI:** 10.1101/2022.04.06.487418

**Authors:** Yuki Kimura, Nobuaki Nakamuta, Masato Nikaido

## Abstract

The evolutionary transition of vertebrates from water to land during the Devonian period was accompanied by major changes in animal respiratory systems in terms of physiology and morphology. Indeed, the fossil record of the early tetrapods has revealed the existence of internal gills, which are vestigial fish-like traits used underwater. However, the fossil record provides only limited data on the process of the evolutionary transition of gills from fish to early tetrapods. This study investigated the gills of *Polypterus senegalus*, a basal ray-finned/amphibious fish which shows many ancestral features of stem Osteichthyes. Based on scanning electron microscopy observations and transcriptome analysis, the existence of motile cilia in the gills was revealed which may create a flow on the gill surface leading to efficient ventilation or remove particles from the surface. Interestingly, these cilia were observed to disappear after rearing in terrestrial or high CO_2_ environments, which mimics the environmental changes in the Devonian period. The cilia re-appeared after being returned to the original aquatic environment. The ability of plastic changes of gills in *Polypterus* revealed in this study may allow them to survive in fluctuating environments, such as shallow swamps. The ancestor of Osteichthyes is expected to have possessed such plasticity in the gills, which may be one of the driving forces behind the transition of vertebrates from water to land.

## Introduction

Fish use their gills for respiration, but these gills progressively degenerated during the evolutionary transition of vertebrates from water to land, in parallel with a corresponding physiological and morphological remodeling of the respiratory system. For example, extant Amniota (mammals, reptiles, and birds) have completely lost their gills and depend on lungs for respiration. In some amphibians, the internal gills which are present during the larval stage degenerate during metamorphosis into adulthood. In tracing back to earlier stages of evolution, the fossil record of Acanthostega, and similar Palaeozoic adult tetrapods, have shown the existence of internal gills (Coates and Clack, 1991; Schoch and Witzmann, 2011), which resembled those of lungfish (Clack, 2012). This suggests that the internal gills were still functioning in the respiratory systems of these early tetrapods which inhabited the interface between water and land. Generally, gills are believed to function only in water because their lamellae collapse due to gravity and dry out in a terrestrial environment (Sayer, 2005). Therefore, the evolutionary degeneration of gills in response to the environmental changes associated with the water to land transition is an issue of interest. Indeed, a few extant species of teleost fish have adapted to terrestrial environments by modifying the structure of their gills (e.g., mudskippers (Low et al., 1988), and mangrove killifish (*Kryptolebias marmoratus*) (Ong et al., 2007)). In particular, plastic structural changes have been observed in the gill lamella of mangrove killifish during acclimation to terrestrial conditions (Ong et al., 2007; Turko et al., 2012).

To better understand the evolution of the respiratory system of stem Osteichthyes, the group *Polypteridae* (bichir, reedfish) provide appropriate model organisms. The *Polypteridae* are the most basal group of living ray-finned fish, and retains several ancestral traits of the stem Osteichthyes. Importantly, some of these traits include the use of lungs or spiracles for air-breathing, and thus they show some of the tetrapod adaptations to survival in a terrestrial environment (Graham et al., 2014; Tatsumi et al., 2016). Some studies argued that the terrestrial adaptation of *Polypterus* occurred independently from that of the tetrapods (Damsgaard et al., 2020; Ord and Cooke, 2016). In the meanwhile, recent genomic, transcriptomic and histological analyses proposed that at least the lungs of *Polypterus* and tetrapods may have originated only once in their common ancestor (Bi et al., 2021; Cupello et al., 2022; Tatsumi et al., 2016). Indeed, Standen *et al*. have successfully kept *Polypterus* on land for up to eight months (Standen et al., 2014). Turko *et al*. found that in *Polypterus* reared on land, cells filled the area between the lamellae of the gills and reduced the size of the gill skeleton (Turko et al., 2019). Therefore, inspection of the gills of *Polypterus* before and after rearing in a terrestrial environment, thus mimicking the water to land transition, may help understand the remodeling process of the respiratory system which occurred during early tetrapod evolution.

In addition to the water to land transition, concentrations of oxygen and carbon dioxide in water also affect gill morphology (Phuong et al., 2017; Turko et al., 2012; Wright and Turko, 2016). Fish excrete carbon dioxide into water through their gills because of the high solubility of carbon dioxide in water (Rahn, 1966); a process which does not work in terrestrial (i.e., atmospheric) or high CO_2_ environments. In fact, *Polypterus* was shown to depress gill ventilation in response to high CO_2_ concentrations in an aquatic environment (Babiker, 1984). Additionally, Ultsch proposed that hypercarbic environments played a role in the water to land transition during vertebrate evolution (Ultsch, 1987; Ultsch, 1996). However, until now, morphological changes to gills upon exposure to such environments have not yet been documented.

In this study, we aimed to gain insight into the evolution of the respiratory system during the water to land transition of the stem Osteichthyes. Specifically, we examined whether changes in gills occur in *Polypterus* after the exposure to high CO_2_ or terrestrial environments, using morphological comparison, immunostaining, and transcriptome as benchmarks.

## Materials and methods

### Animals and rearing environment

The *Polypterus senegalus* Cuvier, 1829, used in this study were obtained from a commercial supplier (Nettaigyo-tsuhan forest, Japan) and kept in normal water conditions for at least one month before being transferred to either terrestrial conditions or high CO_2_ conditions. The total length and the body weight of fish were shown in supplemental table 1. The experiments were conducted in accordance with the Tokyo Institute of Technology Regulations for the Management of Animal Experiments.

The control fish were kept in a 600 L tank with a filtration system. The water temperature was approximately 28°C ± 1°C, with a 12:12 light/dark cycle. Fish in a terrestrial environment were kept in a 25 cm × 35 cm mesh cage at a depth of about 1 mm with a supply of filtered water and foggers to prevent ventral and dorsal surface from drying, respectively (Standen et al., 2014; Turko et al., 2019). In the high CO_2_ experiment, ambient temperature CO_2_ gas was added continuously until the gill ventilation of *Polypterus* was depressed, in accordance with previous research (Babiker, 1984). The concentration of free carbon dioxide in the treatment tank was kept above 100 mg/L (pH 5.08–5.85). The concentration in the control tank was 8.8 mg/L (pH 6.89–7.11). The free carbon dioxide concentration was measured according to the method of the previous study (Sharma, 1998). The 35 cm × 35 cm × 20 cm tank was continuously stirred using a water filter (Rio+ filter-set 2, Kamihata, Japan) to equalize and ensure a uniform carbon dioxide concentration. Fish were kept in the environments described above for at least one month.

### Observation by scanning electron microscopy (SEM)

To examine changes in the microstructure of the surface of the gills, we conducted SEM observation. After keeping the animals in the respective environments, they were euthanized by decapitation and dissected. Specimens of gills for electron microscopy were washed in 0.7x PBS, then fixed in 2.5% glutaraldehyde, and treated with 8N HCl at 60°C for 30 minutes to remove surface mucus. The specimens were then dehydrated by ethanol series, and dried in a freeze dryer (ES-2030, Hitachi, Japan) using t-butyl alcohol. The specimens were osmium coated and observed by SEM (JSM-7001F, JEOL, Japan).

The length of the cilia in the SEM image was measured with Fiji (Schindelin et al., 2012). We selected five representative samples for which the total length could be measured, and calculated the mean and unbiased variance.

### Immunofluorescence staining

To examine the presence/absence of cilia and taste buds in the gills, we conducted immunostaining. Indirect immunofluorescence was performed using the following antibodies: anti-alpha tubulin (acetyl K40, rabbit monoclonal, abcam, ab179484, 1:2000), and anti-calretinin (mouse monoclonal, swant, 6B3, 1:2000) were used as primary antibodies for detecting cilia and taste buds, respectively. Three or more individuals were verified for each condition (Suppelemental table 1). Anti-rabbit IgG (H+L) Alexa Fluor 594 (donkey polyclonal, Invitrogen, AB_141637, 1:2000), and anti-mouse IgG (gamma 1) Alexa Fluor 488 (goat polyclonal, Invitrogen, AB_2535764, 1:2000) were used as secondary antibodies. The entire gills were immunostained as follows: incubated with 0.5 % Triton x-100 (v/v) in 0.7x PBS for 15 minutes at room temperature, washed with 0.7x PBS, and then blocked with normal goat serum 10% and bovine serum albumin 1% in 0.7x PBS for 1 hour. After washing with PBS, primary antibodies of anti-tubulin were added and incubated at room temperature for 1 hour. Secondary antibodies were reacted following the same procedure. The same procedure was also used to stain for calretinin. To prevent fading of the fluorescence, a mounting medium (VECTASHIELD with DAPI, Vector Laboratories, US) was added. The images were taken with a fluorescence microscope (Axioplan2, Carl ZEISS, Germany). After taking the photos, Photoshop was used to correct the levels and modify the colors.

### Histological observation with HE stain

We examined whether the inter-lamellar spaces between the gills of *Polypterus* become filled with cells or not in the terrestrial (Turko et al., 2019) and high CO_2_ environments. Gills specimens were fixed in Bouin’s fixative. The specimens were then dehydrated in ethanol, replaced by xylene, and embedded in paraffin. Thin sagittal sections with a thickness of 5 μm were prepared. The sections were stained with haematoxylin and eosin.

To quantify changes in heights of inter-lamellar cell mass (ILCM) in terrestrial and high CO_2_ environments, we measured the heights of ILCM of individuals in control, terrestrial and a high CO_2_ environment with ImageJ. Two gill lamellae per individual were selected and three ILCM per gill lamellae were randomly selected. To test for changes of heights of ILCM, we compared them using F-test and Welch’s *t*-test with scipy version 1.7.1.

### RNA-seq analysis

To examine the differences in gene expression in gills between aquatic and terrestrial environments, we conducted RNA-seq analysis. RNA was extracted from whole gills of three individuals reared in the control water and in the terrestrial environment using TRI Reagent (Molecular Research Center, Inc., U.S.). The extracted total RNAs were sequenced at 100 bp paired-end on a NovaSeq 6000 from Macrogen Japan Corp, with the TruSeq stranded mRNA Library Prep Kit. The quality control of raw sequence data was performed with fastp (Chen et al., 2018) with the following options: -g, -q 20, -w 16. Next, the read data were mapped to the reference genome of *Polypterus senegalus* (Bchr_013 (Bi et al., 2021)) with STAR version 2.7.3a (Dobin et al., 2013). The mapped reads were then counted with featureCounts (Liao et al., 2014). Differentially expressed genes (DEGs) were estimated by TCC (Sun et al., 2013) with the iDEGES/*edgeR*-*edgeR* combination (p < .05). The Over Representation Analysis (ORA) was conducted with WebGestalt (http://www.webgestalt.org) (Liao et al., 2019).

### Recording gill ventilation and water flow

We recorded the head movement of the *Polypterus* to see the difference in gill ventilation behavior between terrestrial and aquatic environments. In the terrestrial environment, the *Polypterus* was placed on a damp non-woven cloth. A digital camera X100V (Fujifilm, Japan) was used for recording.

To detect water flow on the surface of the gills, *Polypterus* blood cells were dropped onto the surface of the excised gills of *Polypterus*. The blood cells were collected at the time of decapitation. The size of red blood cells of *Polypterus palmas* is 160 μm^2^ (Hardie and Hebert, 2003). Before blood cell drops, we confirmed that there was no bleeding from the gills themselves. The movement of blood cells was recorded using microscopes (Axioplan2, Carl ZEISS, Germany and SZX7-TR2-APO-C, OLYMPUS, Japan). The movies were edited at 10x and 4x speed respectively.

## Results

### The cilia on the surface of gills

SEM observations and immunostaining with an acetylated tubulin antibody revealed the existence of cilia as bundle-like structures distributed on the surface of primary gill lamella of *Polypterus senegalus* (gray bichir) (Fig. 1A–C and Fig. 2A). The mean length of the cilia was 3.93 μm and unbiased variance was 0.16. The cilia were distributed in a dotted pattern.

**Figure 1.**
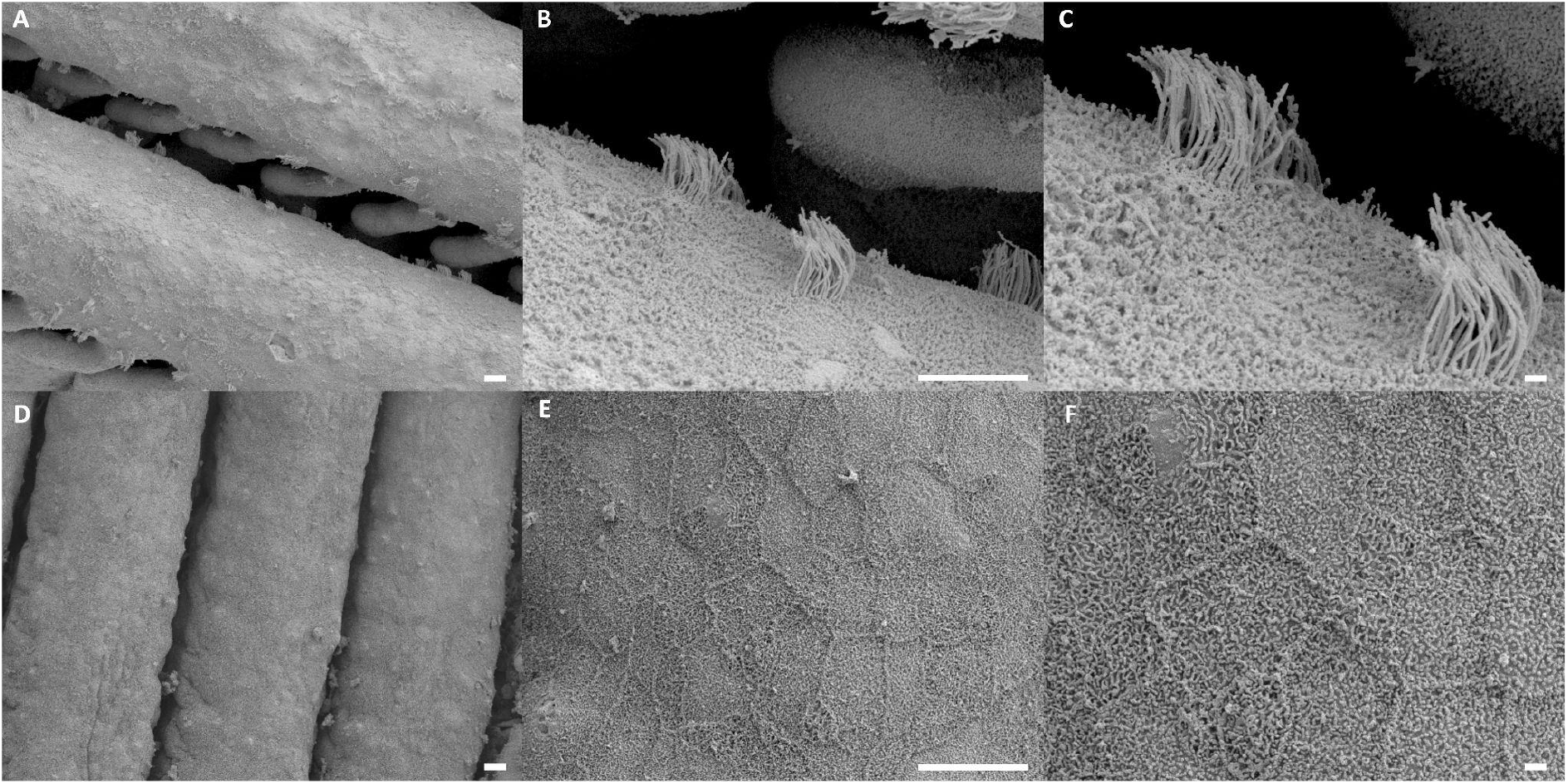
Scanning electron micrograph of the surface of gill filaments of *P. senegalus*. The surface of *Polypterus* reared in water (A–C) and terrestrial environment (D–F). Scale bar in A, B, D and E = 10 μm, in C and F = 1 μm.

**Figure 2.**
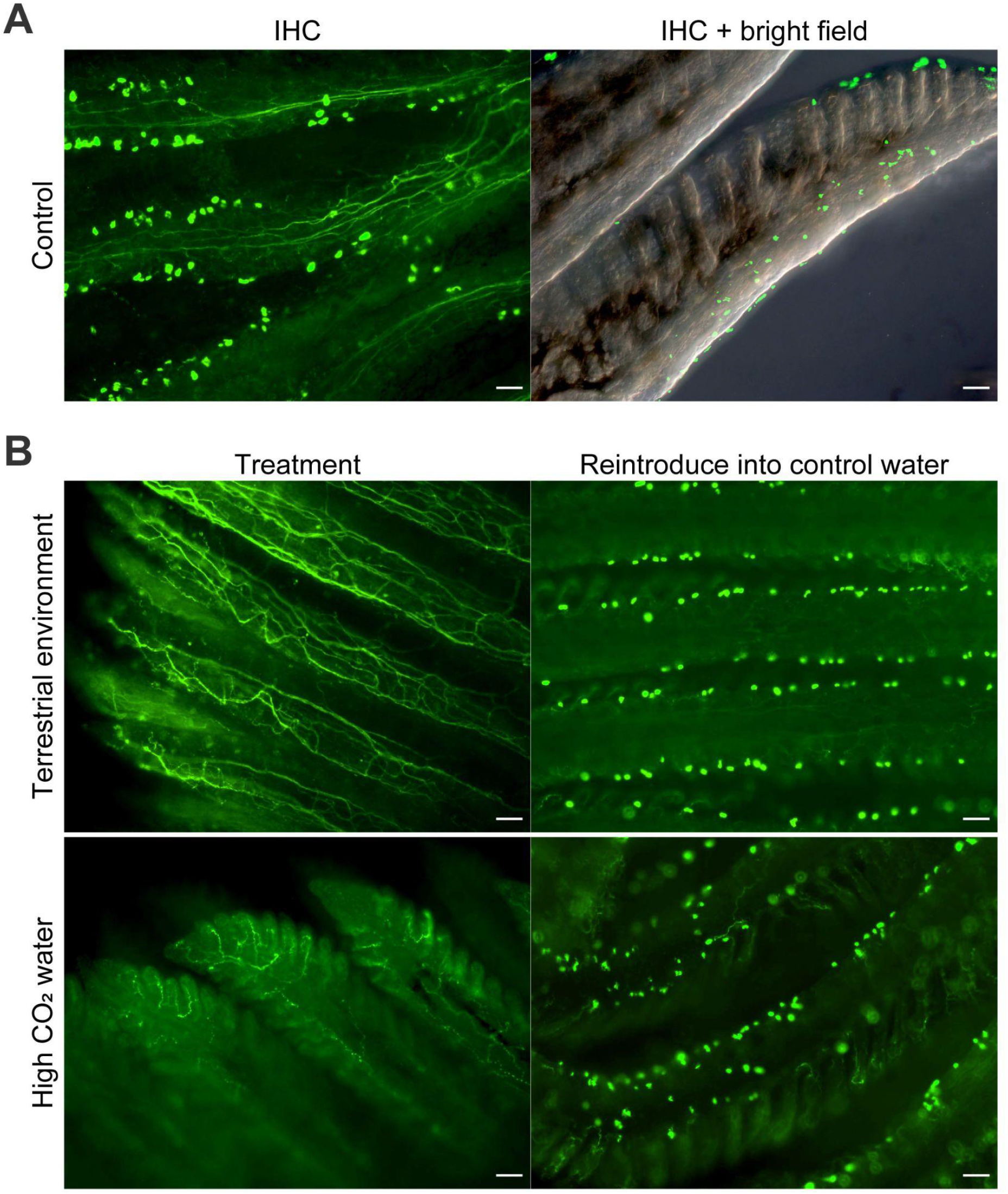
Acetyl alpha tubulin immunostaining of gills of *P. senegalus*. The gills of *Polypterus* reared in control water (A), in a terrestrial environment (B: top part), and in high CO_2_ water (B: bottom part). Punctate spots indicate the presence of multiple cilia. Scale bar = 50 μm.

Next, individual *Polypterus* were reared in either terrestrial environments or in high CO_2_ water for more than one month. It is noteworthy that gill ventilation was observed when the water level was high enough to immerse the head, but was suppressed when the water level was too low to immerse the head (Supplemental movie 1). The suppression of gill movement was also observed in the high CO_2_ environment, as was also observed by (Babiker, 1984). The gills of *Polypterus* reared in terrestrial and in high CO_2_ water environments were then investigated to identify the presence or absence of cilia. Interestingly, SEM and immunostaining observation revealed that the cilia disappeared in the gills of *Polypterus* reared in both of these environmental conditions (Fig. 1D–F, Fig. 2B).

In the next trial, we returned the *Polypterus* individuals exposed to terrestrial or high CO_2_ environments for one month back to the original water environment and reared them for an additional month. The immunostaining observations for the resultant *Polypterus* individuals revealed that the cilia in the gills had recovered (Fig. 2B). These results suggest that the *Polypterus* possesses the ability to undergo plastic morphological changes (formation and de-formation) of cilia in the gills in response to environmental changes they experience.

In addition to the gill cilia, the enlargement of the inter-lamellar cell mass (ILCM) between the lamellae of the gills was examined in the *Polypterus*, which were reared in both the terrestrial and high CO_2_ environments. A previous study revealed that enlargement occurs in the ILCM of *Polypterus* reared on land (Turko et al., 2019), which was confirmed in the present study (Fig. 3). In addition, a statistically significant enlargement of the ILCM was also observed in the individuals reared in the high CO_2_ environment (Fig. 3).

**Figure 3.**
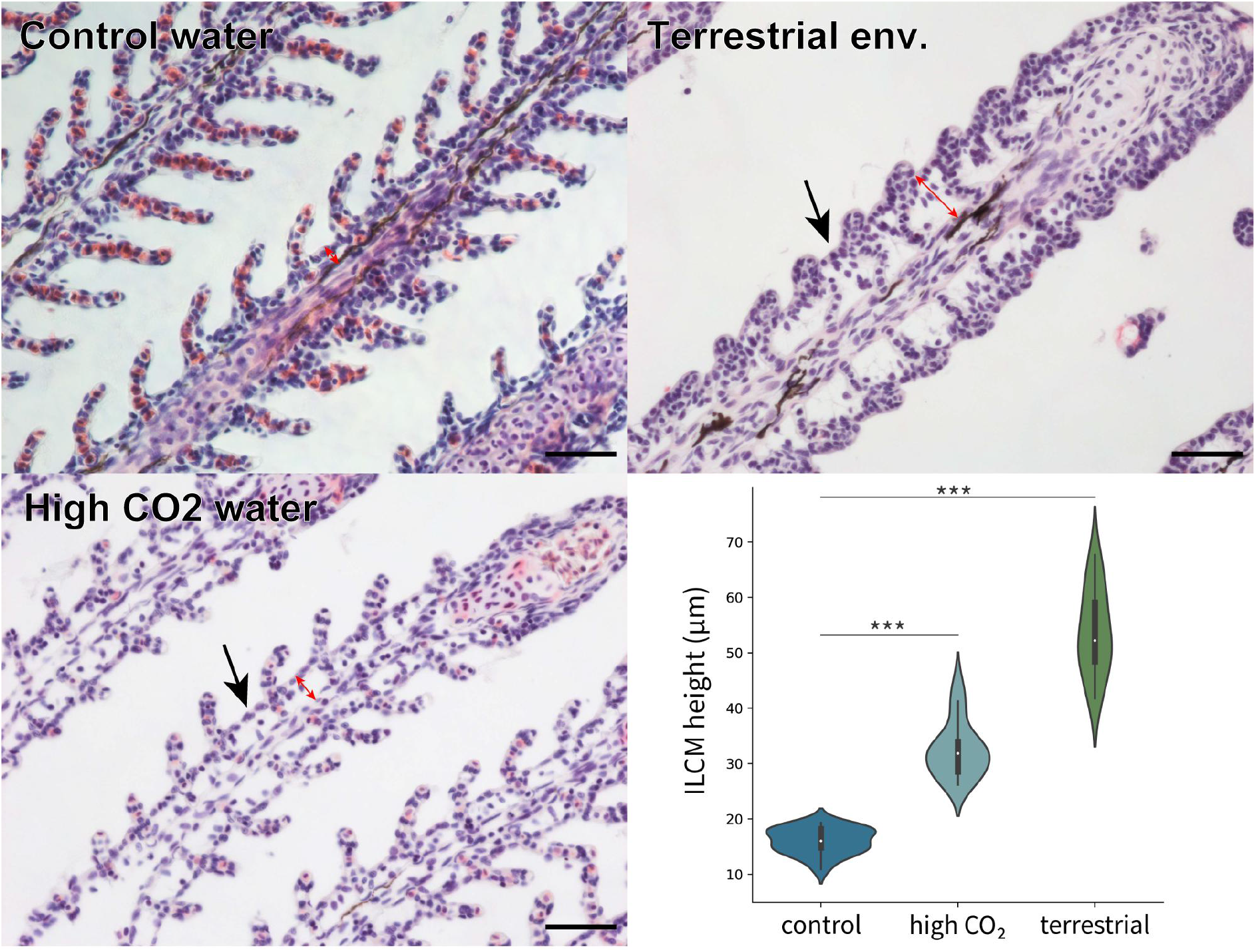
Light micrographs of gills of *P. senegalus* stained with hematoxylin and eosin. The inter-lamellar cell mass (ILCM) was enlarged in high CO_2_ water condition and terrestrial environment compared to the control water (arrows). The red arrows indicate the height of ILCMs. Scale bar = 50 μm. The ILCM height of individuals in control, high CO_2_, and terrestrial conditions were measured. Welch’s *t*-test was calculated. ***: P-values significant at < 0.0001.

### Elucidation of cilia function by RNA-seq analysis

There are many types of cilia, which are classified into two main groups: motile and non-motile cilia. They are various in terms of their locations (e.g. airways, olfactory epithelium) and functions (Choksi et al., 2014). We next investigated which type of cilia we found in the gills of *Polypterus*.

The DEGs analyses were conducted on the whole exome sequencing data (RNA-seq data) for the gills of the *Polypterus* reared in both water and terrestrial environments. In particular, this study aimed to elucidate the function of cilia in the gills, which had not yet been documented. Considering that cilia were lost in the terrestrial environments, a focus was made on the DEGs of which the expressions were reduced in gills under terrestrial conditions. As a result, a total of 868 DEGs were obtained which was statistically significant. An ORA was then performed for the list of down-regulated DEGs (p < .05) using WebGestalt (Liao et al., 2019). The results of the ORA indicated that the down-regulated gene groups were related to 1. axonemal dynein complex assembly, 2. axoneme assembly, 3. cilium movement, and 4. cilium assembly (Fig. 4A). The further inspection of these gene groups revealed that they were represented by *foxj1a*, which is included in one of the major genes involved in cilium movement (Fig. 4B). *foxj1a* is a homolog of *foxj1* (Forkhead box protein J1)(Aamar and Dawid, 2008; Hellman et al., 2010). The protein encoded by *foxj1* is a master regulator of the motile ciliogenic program (Yu et al., 2008). The degree of expression of several genes, known to be regulated by *foxj1* (Mukherjee et al., 2019) was examined, and found that they were also significantly down-regulated (Fig. 4B). These results indicate that *foxj1*, and its downstream genes involved in cilia movement, were down-regulated in terrestrial environments. These lines of data suggest that the cilia distributed on the gills of *Polypterus* are motile.

**Figure 4.**
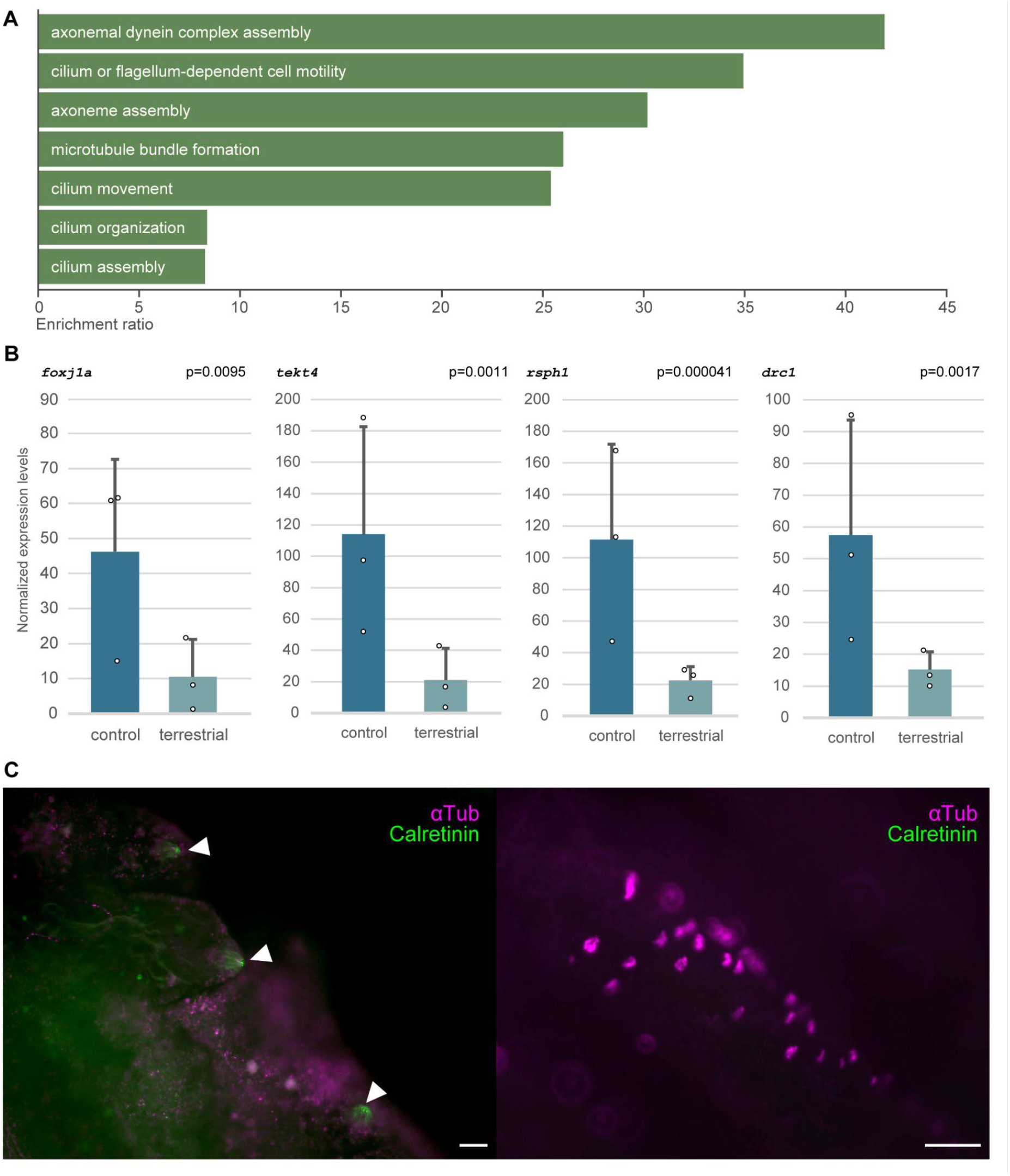
Results of expression analysis and calretinin immunostaining in gills. (A) The results of over representation analysis using genes whose expression levels were reduced in the gills of *Polypterus* reared in a terrestrial environment. (B) Normalized expression levels of four representatives of genes known to be involved in cilia movement. TCC was used for normalization (see methods section). The dots on the plots indicate individual expression levels. (C) Micrograph of gills immunostained with anti-calretinin antibody (light green) and anti-acetylated alpha tubulin antibody (magenta). The left image shows the presence of taste buds (arrowhead) and absence of ciliated cells in the gill arches. The right image shows the presence of ciliated cells and absence of taste buds in the gill lamella. Scale bar = 50 μm.

To confirm whether the cilia were motile, we observed the water flows generated on the surface of the gills. The blood cells of *Polypterus* were used for visualizing the flow. The movement of blood cells was observed from the root to the tip of the gill lamella (Supplemental movie 2). The water flow was also observed in the movie of the entire gill (Supplemental movie 2), confirming that these cilia are motile.

Whether these cilia have specific functions beyond motility was also examined. It has been shown that taste buds of lampreys possess cilia, while those of bony fishes and mammals possess microvilli (Baatrup, 1983; Barreiro-Iglesias et al., 2008). However, until now, there have been no reports describing the taste buds of *Polypterus* in terms of the existence of cilia or microvilli. Therefore, an examination was conducted to determine whether the ciliated cells on the gills of *Polypterus* were taste buds or not. Immunostaining was performed using an anti-calretinin antibody which is used as a chemosensory marker in vertebrates, including for taste buds (Barreiro-Iglesias et al., 2008). As a result, clear signals of calretinin-positive cells were observed which show typical taste bud like structures on the gill arches of *Polypterus* (Fig. 4C left). Importantly, no acetylated tubulin-positive signal was observed, indicating the absence of ciliated cells in the taste buds. Additionally, no calretinin-positive cells were observed where acetylated tubulin positive cells were concentrated (Fig. 4C right). These results suggest that the ciliated cells on the gills of *Polypterus* are not taste buds.

Whether these cilia possessed functions of innate immunity was also examined, since the ciliated cells found in the airway epithelium of mammals have been shown to be involved in the immune system (Freund et al., 2018; Shah et al., 2009). Indeed, several taste 1 receptors (*Tas1R*s) and taste 2 receptors (*Tas2R*s) expressed in the upper airway respond to substances secreted by Gram-negative bacteria. These *Tas1Rs* and *Tas2Rs* are essential to trigger mucociliary clearance by controlling the beating of motile cilia (Lee et al., 2012). Therefore, the expression of the *Tas1R*s and *Tas2R*s gene cascade was examined. The expression of *Tas1Rs* were not changed significantly (Fig. S1A). The expression level of *Tas2R*s was quite low and difficult to detect by RNA-seq. The next focus was on *trpm5* and *plcb2*, which are located downstream of the *Tas1R*s and *Tas2R*s gene cascades (Ahmad and Dalziel, 2020; Tuzim and Korolczuk, 2021). These genes were expressed in gills, but there were no significant changes in expression levels between aquatic and terrestrial environments (Fig. S1B). Next, the focus was on three genes of nitric oxide synthase (NOS). Nitric oxide controls the beating of the cilia in the airway epithelium of mammals (Li et al., 2000). It is noteworthy that one of the NOS genes was lost in teleost fish after the whole genome duplication (Donald et al., 2015), however, all three NOS genes were identified in the genome of *Polypterus*. A comparison of the expression levels of the three types of NOS genes in the gills of *Polypterus* found no significant differences between the control and terrestrial environments (Fig. S1C). These lines of data suggest that the cilia in the gills of *Polypterus* do not play a specific role in Tas1R/Tas2R-related innate immunity shown in the mammalian airway epithelium, but provide only motile functions, which may contribute toward producing water flow for efficient ventilation.

In addition to the cilia function, we examined the possibility of whether or not the loss of cilia in the terrestrial environment was the result of injury. We inspected the up-regulated DEGs which contain 1,473 genes, showing that the genes associated with injury (e.g. inflammation) were not enriched. On the other hand, the genes associated with migration, localization, and development were enriched (Fig. S2).

## Discussion

### Origin of cilia in the gills

This study revealed the existence of cilia in the internal gills of *Polypterus*. This is the first report of ciliated cells of the internal gills in a ray-finned fish (Actinopterygii). Although several SEM images of gills have been presented in model organisms such as zebrafish (*Danio rerio*) and medaka (*Oryzias latipes*) (Leguen, 2018; Messerli et al., 2020), the cilia were not described in these studies. Cilia were not found in gills and/or specialized respiratory organs in walking catfish (Clariidae) nor in mangrove killifish (*K. marmoratus*), which are also adapted to the terrestrial environment in a manner similar to *Polypterus* (Maina, 2018; Ong et al., 2007). Additionally, sharks (cartilaginous fishes) do not possess cilia in their gills (Bullard et al., 2001). On the other hand, cilia have been reported in the internal gills of a lobe-finned fish, the Australian lungfish (*Neoceratodus forsteri*, Sarcopterygii) (Kemp, 1996), which diverged from the common ancestor of Osteichthyes 470 million years ago (Wang et al., 2021). It is plausible that cilia in the internal gills had been acquired in the common ancestor of Osteichthyes and retained in *Polypterus* and lobe-finned fish, but were lost in most other ray-finned fish, which diverged later than *Polypterus*. However, the size and arrangement of the cilia in the gills of *Polypterus* are distinct from those seen in lungfish (Kemp, 1996). Indeed, the cilia of *Polypterus* are noticeably short and are distributed in a circular dotted pattern (Fig. 1), while those in lungfish are long and distributed in band-like patterns (Kemp, 1996). The morphological distinctness of the cilia observed in the gills of *Polypterus* and lungfish implies that specific roles of cilia have diversified between ray-finned and lobe-finned fish.

### Function of the cilia

In this study, immunostaining data showed that the cilia found in the internal gills of *Polypterus* were not taste buds. Additionally, transcriptome analyses suggested that these cilia did not possess functions in Tas1R/Tas2R related to innate immunity as seen in the airway epithelium of mammals. On the contrary, the cilia seen in the internal gills of Australian lungfish, which exist only during the larval stages, have been shown experimentally to produce water flow (Kemp, 1996). These cilia are responsible for increasing the efficiency of ventilation and eliminating small particles such as microorganisms from gill surfaces (Kemp, 1996). In this study, the cilia found in the gills of adult *Polypterus* were shown to be motile (Fig. 4, Supplemental movie 2). The roles of the cilia may be explained by the dual ventilation system used by *Polypterus*, which utilizes both lungs and gills. A previous study revealed that the frequency of gill ventilation in *Polypterus* decreases significantly before and after air-breathing (Magid, 1966). *Polypterus* may reduce its frequency of gill ventilation by complementing it with pulmonary ventilation. However, depression of gill ventilation leads to the reduction of other gill functions such as eliminating small particles, ion transport, excretion of carbon dioxide and ammonia (Ultsch, 1996). During such temporary depression of gill ventilation, the motile cilia may function to increase efficient water flow for excretion and transport. The constant direction of water flow produced by the cilia also reinforces the hypothesis (Supplemental movie 2). Considering that gill cilia and pulmonary ventilation are expected to be mutually reinforcing in function, it is plausible to speculate that they were acquired simultaneously in their evolution.

### The plasticity of cilia in gills in response to the environments

This study found that the cilia in the internal gills of *Polypterus* were lost after the exposure to a terrestrial environment (Fig. 1, 2). Assuming that cilia play an essential role in efficient ventilation, excretion of carbon dioxide, and removal of small particles, it makes sense that they would disappear under conditions where water is unavailable. It is also likely that the loss of cilia may play a role of saving consumption of energy in a terrestrial environment where feeding is restricted. Previous studies have revealed a reduction in the volume of mineralized bone in the gills after 8 months of rearing on land, arguing that *Polypterus* may reduce investment in its gills when in a terrestrial environment (Turko et al., 2019). Importantly, the loss and regeneration of the cilia are plastic in response to the environmental change (Fig. 2). Given that no experimental data showing the signal of inflammation was obtained (Fig. S2), the plastic loss and regeneration of cilia may be programmed biological responses rather than just an injury. This plasticity may have been acquired as a result of an adaptation to fluctuating environments, such as encountered in shallow rivers and swamps, which *Polypterus*, and possibly the ancestors of Osteichthyes inhabited.

Similar to terrestrialization, the high concentration of CO_2_ in water resulted in the plastic loss and regeneration of cilia in the gills of *Polypterus* (Fig. 2). Previous study has revealed that gill ventilation in *Polypterus* was depressed in water with high CO_2_ concentrations (Babiker, 1984). In a high CO_2_ environment, gill ventilation causes several negative impacts, represented by acidosis. Since CO_2_ relies on passive transport, elevated CO_2_ concentrations in environmental water may lead to an influx of CO_2_ into the blood through the gills. Hence, suppressing gill ventilation and reducing exposure of the gills to high CO_2_ water is expected to be adaptive in *Polypterus*. Apart from the gills, the kidney and intestine are also able to exchange ammonia and ions. Therefore, even while gill ventilation is depressed, those organs may compensate for this function. The CO_2_ concentration in the water tends to transiently increase in warm, relatively shallow and vegetated environments. The Devonian air-breathing vertebrates are considered to have inhabited environments with fluctuating CO_2_ concentrations (Ultsch, 1996). *Polypterus* also inhabits similar environments (Magid, 1967). It is presumed that *Polypterus* survives in a high CO_2_ environment by depressing gill ventilation and switching to ventilation through the lungs when needed. To prevent exposure of the gills to high CO_2_ water, the frequency of gill ventilation as well as the motile cilia, would likely be reduced.

### ILCM size changes in response to environmental changes

In addition to these microstructural changes involving cilia, the enlargement of the ILCM was observed in response to high CO_2_ concentrations or when the animal was in terrestrial environments (Fig. 3). A previous study showed that the ILCM between the secondary gill lamellae were enlarged in the gills of *Polypterus* reared on land (Turko et al., 2019). In this study, we found that the genes associated with tissue development and cell migration were upregulated in a terrestrial environment (Fig. S2). The upregulation of these genes were consistent with the enlargement of the ILCM. The enlargement or reduction of ILCM are also observed in teleost fish such as mangrove killifish under terrestrial conditions (Ong et al., 2007). The enlargement of the ILCM of *Polypterus* in high CO_2_ conditions suggests a reduction of gas exchange surface area of the gills, leading to a presumed reduction of the passive import of CO_2_ from the water. It is noteworthy that both the high CO_2_ condition and terrestrial environment induced the similar morphological changes in the gills of *Polypterus*.

### High CO_2_ environments and the water to land transition

In previous studies, the possible link between hypercarbia and the water to land transition has been discussed. Because of its high solubility in water, CO_2_ is normally excreted through the gills of most fish. In contrast, air-breathing fish take in O_2_-rich air, resulting in a lower frequency of ventilation using gills. Therefore, the blood of air-breathing fish shows a higher *P*_CO2_ relative to other fish (Bayley et al., 2019; Ultsch, 1996). Indeed, in some obligate air-breathing fish, the acid-base status of the blood is similar to that seen in amniotes (Bayley et al., 2019; Ultsch, 1996). Based on studies showing that hypoxic environments are also high in carbon dioxide, it has been proposed that air-breathing fish may have shifted to high *P*_CO2_, [HCO_3_^-^] blood prior to their terrestrial adaptation during vertebrate evolution (Ultsch, 1996).

It is noteworthy that both high CO_2_ water and terrestrial environments were found to lead to the similar results in *Polypterus*, namely, the depression of gill ventilation, ILCM enlargement, and the loss of cilia in the gills. Based on the present and previous findings (Bayley et al., 2019; Ultsch, 1996), we support that adaptive traits to a high CO_2_ environment shown above were acquired prior to terrestrial adaptation. Namely, the ancestors of stem tetrapods possibly become ready to adapt to land because some adaptive characters acquired in high CO_2_ environments were also adaptive in terrestrial environments. For example, the depression of internal gill ventilation and enlargement of ILCM was adaptive in terrestrial environments in terms of preventing desiccation. The loss of cilia was also adaptive in that respiration and material transfer through internal gills are not functional on land. Importantly, above changes of internal gills were plastic in the ancestor of Osteichthyes, that may allow them to accomplish the drastic transition from water to land back and forward, and step by step.

In this study, SEM and immunostaining observations and transcriptome analyses revealed the existence of motile cilia in the internal gills of *P. senegalus*, which may create a flow of water on the gill surface leading to efficient respiration and material transport. These motile cilia were observed to have been plastically lost and regenerated in response to environmental changes resulting from rearing in terrestrial and high CO_2_ environments. In some Devonian fish, some characters evolved under similar high CO_2_ environments may have been later co-opted during the adaptation to terrestrial environments by early tetrapods. The specific features characterized in the present study of *Polypterus*, the oldest lineage of existing amphibious ray-finned fish, provide important insights into the evolutionary transition of vertebrates from water to land.

## Acknowledgements

We thank Shizuka Oki, Yasuyo Shigetani, and Masataka Okabe of the Jikei University School of Medicine for teaching HE staining.

## Data availability

All sequence reads were deposited in the DDBJ Sequence Read Archive under BioProject accession no. PRJDB13172.

## Authors’ contributions

Y.K.: Conceptualization, formal analysis, investigation, writing - original draft, visualization, funding acquisition; N.N.: Investigation, writing - review & editing; M.N.: Writing - review & editing, supervision, funding acquisition

## Figures

**Figure S1.**
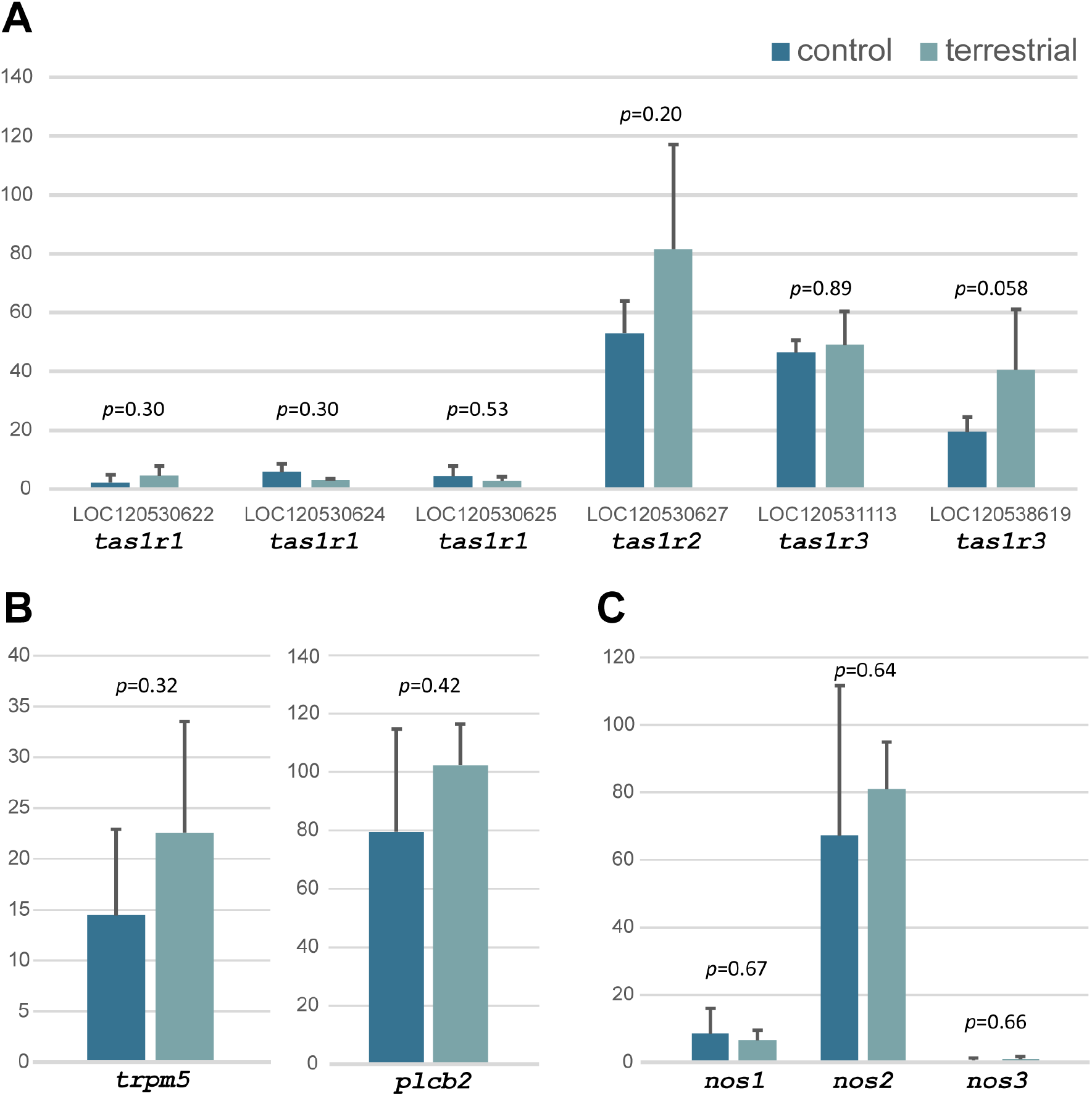
Expression levels of some genes in gills in aquatic and terrestrial environments. Normalized expression levels of (A) Tas1Rs and genes known to be involved in (B) Tas2R cascade (C) NOS (Nitric Oxide Synthase). TCC was used for normalization and testing (see methods section).

**Figure. S2.**
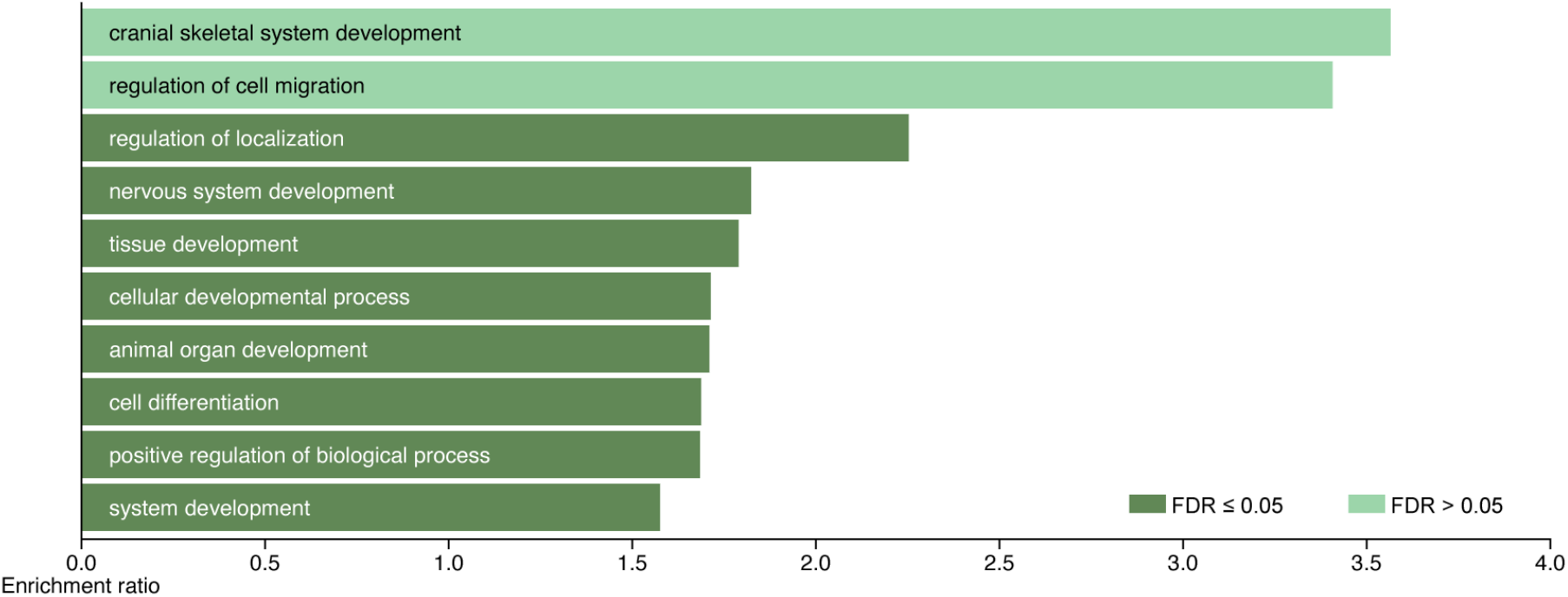
The results of over representation analysis for up-regulated genes in a terrestrial environment. The genes that were significantly upregulated were analyzed (see methods section).

**Supplemental table 1. Profile of individuals used in the experiments**.

**Supplemental movie 1. Gill ventilation of *Polypterus* in water and terrestrial environments**. The gill ventilation of *Polypterus* in water (left side) and terrestrial environments (right side). Both movies are the same speed.

**Supplemental movie 2. The water flows on the surface of the gill**. The blood cells of *Polypterus* visualize water flow on the surface of the gill lamellae. In the first half of the video, the water flow is generated from the root (right side) to the tip (left side) of the gill lamellae, and in the second half of the video, the water flow is generated from the root (underside) to the tip (upperside). The movies were edited at 10x (first half) and 4x (second half) speed respectively.

